# PhyTaxR, a set of phytoplankton taxonomic curation tools in R

**DOI:** 10.64898/2026.10.23.761320

**Authors:** Fernando Rayón-Viña, Zoe V. Finkel, Andrew J. Irwin

## Abstract

Taxonomic harmonisation is a persistent challenge for phytoplankton monitoring and biodiversity synthesis. Operational taxon fields often combine scientific names —some of which may be incomplete, misspelled, or otherwise non-standard— with size classes, morphological descriptors, uncertainty qualifiers, vernacular labels, author citations, and historical nomenclature. Here we introduce PhyTaxR, an R package that formalizes the curation of phytoplankton taxonomic records as a reproducible, modular workflow. The package separates a standardized taxonomic core from ancillary information while preserving uncertainty and descriptive metadata. It then applies a conservative taxonomic-resolution workflow based primarily on searches of the World Register of Marine Species (WoRMS), supplemented where appropriate by stringent, cross-validated matches from Global Biodiversity Information Facility (GBIF). Matched records are reconciled to accepted taxonomic concepts and returned with persistent identifiers, resolution provenance, and a tidy taxonomic hierarchy. Labels that cannot be resolved automatically are directed to a semi-automatic review workflow, where users can evaluate candidate matches, supply manual identifications, flag records for expert review, or remove unsuitable entries. Illustrative applications show how ‘PhyTaxR’ can harmonize synonyms and spelling variants, retain contextual information associated with original labels, and make unresolved or uncertain records explicit rather than forcing unsupported assignments. By retaining intermediate decisions and supporting targeted expert intervention, ‘PhyTaxR’ reduces manual curation while keeping taxonomic uncertainty visible in reproducible R workflows.

## Introduction

Biodiversity research increasingly relies on the integration of large datasets assembled from multiple sources, time periods, and taxonomic authorities [1, 2, 3]. Taxon names provide the link between individual observations and the biological entities to which they are assigned, and are therefore essential for combining, comparing, and interpreting records across studies. However, this link is often unstable: the same taxon may be recorded under alternative spellings, synonyms, or historical names, whereas changes in taxonomic concepts and classifications can alter the relationship between a name and the entity it denotes. These difficulties are particularly consequential when data span decades, because historical identifications may no longer correspond directly to current taxonomic usage. If such changes are not documented and reconciled, records of the same biological entity can be retained under multiple names, or observations assigned to different taxonomic concepts can be inadvertently combined, potentially biasing estimates of biodiversity patterns, long-term trends, and cross-study comparisons. Taxonomic name harmonisation—the process of linking recorded names to explicit, curated taxonomic concepts while retaining information on synonymy and taxonomic change—is therefore a necessary step in biodiversity data integration rather than a purely cosmetic standardisation procedure [4, 5, 6].

This situation has led to the development of a growing ecosystem of taxonomic databases and tools designed to address these challenges, including global backbones such as the GBIF Backbone Taxonomy [7] and Catalogue of Life [8], as well as domain-specific resources such as the World Register of Marine Species (WoRMS) [9] for marine taxa. In parallel, several R packages provide programmatic access to taxonomic services and name-resolution workflows, including general-purpose tools such as taxize [10] or the Taxonomic Name Resolution Service (TNRS/RTNRS) [11]. However, most of these tools are designed as broad, taxon-agnostic interfaces to online databases and typically assume relatively clean scientific names as input. They rarely account for the idiosyncratic, “noisy” labels found in operational monitoring datasets, where taxon fields often combine scientific names with morphological descriptors, size classes, uncertainty markers, vernacular terms, and observer-specific abbreviations. This gap is particularly acute for marine and estuarine phytoplankton monitoring, where routine counts may include entries such as “unidentified diatom 10–20 µm”, “gymnodinioid cysts”, or “green flagellate, small”. Consequently, many phytoplankton observatories and databases have relied on bespoke, semi-manual curation scripts that implement complex heuristic cleaning and name-resolution rules. Recent qualitative research has highlighted the fragmentation of software practices in marine science, where researchers often learn tools informally, develop individualized or ad hoc workflows, and may encounter barriers to discovering, evaluating, and reusing existing tools [12]. Although such scripts may be effective in their original context, they are often difficult to reuse, test, and maintain, and they rarely expose their decision rules in a transparent and reproducible form. There is therefore a need to formalize these workflows as well-documented, modular tools that integrate seamlessly with modern data-analysis ecosystems.

Here we introduce PhyTaxR [13], an R package for the curation and identification of phytoplankton taxonomic names through a reproducible workflow for cleaning and harmonization. The package combines three sequential steps: (1) a dedicated cleaning pipeline that separates the taxonomic “core” of each entry from ancillary descriptors, recorded as epithets; (2) a conservative automatic resolution workflow that derives accepted names and identifiers through exact and Taxamatch-based searches [14] in WoRMS, together with high-confidence matches from GBIF cross-validated against WoRMS where possible; and (3) a user-mediated fuzzy resolution stage for the most ambiguous cases. The cleaning step is central to this design because real-world taxon fields often combine scientific names with size ranges, morphological descriptors, uncertainty markers, author citations, informal group labels, and spelling or typographical errors in otherwise valid Latin names. Such noisy entries may prevent direct matching against taxonomic backbones. PhyTaxR therefore applies modular text-processing steps to normalize character encodings, extract and preserve ancillary information, standardize infraspecific rank notation, and separate vernacular or generic labels from the taxonomic core before attempting resolution. By transforming heterogeneous taxon labels into standardized, taxonomically interpretable inputs, the cleaning stage substantially improves the performance of the subsequent conservative automatic resolution workflow. In the illustrative application presented here, the cleaning-enabled workflow automatically resolved 47 of the 52 records produced by the cleaning stage

(90.4%), leaving five records for user-mediated review in Stage 3.

Cleaned names are subsequently evaluated through conservative automatic matching, including similarity-based searches that can identify likely misspellings while accepting only matches that meet predefined confidence criteria. The remaining ambiguous entries are then handled through explicit user review in the fuzzy-matching stage.

PhyTaxR is implemented in R and designed to integrate naturally with the modern tidyverse syntax and workflows. The package provides both low-level functions for each cleaning and resolution step and high-level wrappers that operate on tidy data frames, making it straightforward to embed taxonomic curation pipelines into reproducible scripts, notebooks, or automated workflows. In this paper, we describe the architecture of PhyTaxR, detail the cleaning and resolution pipelines, and illustrate their application to phytoplankton monitoring data. We then discuss the strengths and limitations of this approach and outline future extensions, including broader taxonomic coverage and integration with additional data standards.

## Materials and Methods

### Software implementation, requirements, and availability

All analyses were conducted using the version 0.3.2 of the package PhyTaxR [13]. The package requires R version 4.1.0 or later, is distributed under the MIT license, and the version used in this study is available through the archived GitHub release tag v0.3.2, which can be installed using devtools::install_github(“RayonVina/phytaxr@v0.3.2”).

The package relies on several R libraries for data manipulation, text processing, access to taxonomic services, and parallel computation. Data manipulation and workflow operations use several packages from the tidyverse, including `dplyr, tibble, tidyr`, and purrr [15]. Text normalization and string comparison use `stringr, stringi`, and stringdist; communication with external taxonomic services relies on worrms [16], httr, and jsonlite; and parallelized operations use `future` and `furrr`. Additional dependencies, including R.utils and progress, support timeout handling and progress reporting.

### Input data and output structure

The minimum input to the cleaning workflow is a data frame containing a character column of original taxonomic labels, conventionally named `taxon`. This is the default source column used by `run_cleaning_pipeline()`, although users can specify an alternative existing column through the col argument (col = “taxon”). The workflow can also accept a character vector of taxon labels, which is internally converted to a data frame. The original input column is retained untouched throughout the workflow.

During the cleaning stage, `PhyTaxR` generates three core fields: `taxon_clean`, containing the standardized taxonomic component used for name matching; `tax_epithet`, containing ancillary information extracted from the original label; and `uncertain`, a logical flag that identifies records containing qualifiers or other indications of taxonomic uncertainty. Internal schema-checking functions ensure that these fields are available when individual cleaning functions are used independently.

The automatic resolution workflow adds fields recording the matched and accepted taxonomic names, identifiers, taxonomic status, resolution method, and explanatory notes. WoRMS assigns each taxonomic name record a unique numeric identifier, termed an AphiaID, which provides a direct link to its taxonomic information. The output fields include `matched_name, accepted_name, matched_aphiaid, accepted_aphiaid, taxonomic_status, resolution_method`, and `resolution_notes; matched_aphiaid` identifies the matched WoRMS record, whereas accepted_aphiaid identifies the currently accepted record to which it is linked. For records resolved through WoRMS, the package can also retrieve a taxonomic classification from kingdom to forma. The high-level workflow, input requirements, and principal data products of the three stages are summarized in Table 1. Supplementary Table S1 provides a complete function-level reference, including individual functions, arguments, default values, and links to the versioned source code.

**Table 1.** High-level PhyTaxR workflow, input requirements, and principal data products.

| Stage | High-level function | Main input | Principal data products and role |
| --- | --- | --- | --- |
| Input data | None | A character vector of raw taxonomic labels, or a data frame containing a character source column. The default column is `taxon`; an alternative can be specified through `col`. | Retains the original taxonomic label to preserve traceability throughout the workflow. |
| Stage 1: Cleaning | `run_cleaning_pipeline()` | Raw taxonomic labels stored in a data frame. Alternative input-column names can be supplied where supported. | Creates `taxon_clean`, containing the standardized taxonomic component used for matching; `tax_epithet`, containing extracted ancillary information; and `uncertain`, indicating taxonomic uncertainty. |
| Stage 2: Automatic resolution | `run_resolution_pipeline()` | A cleaned data frame containing at least `taxon_clean`; normally the output of Stage 1. | Adds matched and accepted names, AphiaIDs, taxonomic status, resolution method, explanatory notes, and, when available, the WoRMS taxonomic hierarchy from kingdom to forma. |
| Stage 3: Semi-automatic fuzzy resolution | `process_fuzzy_batch()` | Records unresolved after Stage 2, together with resolution fields and optional genus and epithet vocabularies. | Generates and presents candidate matches for explicit user review. Decisions can be accepted, retained for expert review, or flagged for removal; review progress can be saved and resumed. |

### Workflow overview

PhyTaxR implements a three-stage workflow: (1) taxon-name cleaning, (2) conservative automatic taxonomic resolution, and (3) semi-automatic fuzzy resolution of the remaining ambiguous records (Fig 1). Each stage retains the original record and stores intermediate or final decisions in explicit output fields, allowing users to inspect the provenance of each taxonomic assignment.

**Figure 1.**
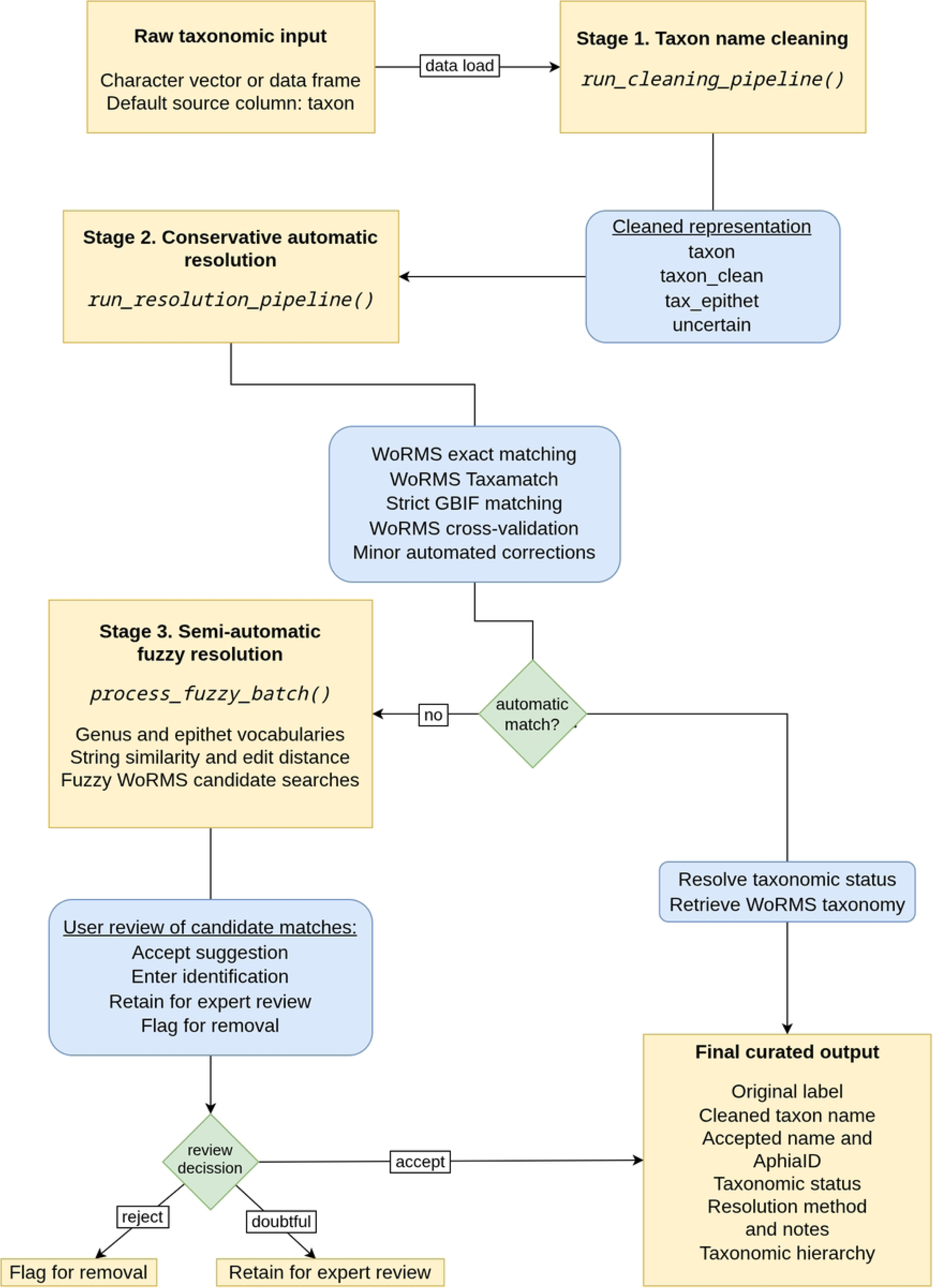
Overview of the three-stage taxonomic-curation workflow implemented in PhyTaxR. Raw taxonomic labels are cleaned to produce a standardized taxonomic representation, then processed through conservative automatic resolution using WoRMS and GBIF. Remaining unresolved records undergo semi-automatic fuzzy matching and user review. The workflow retains intermediate fields and resolution metadata to support traceability.

### Stage 1: Taxon-name cleaning

#### Schema initialization and character normalization

The cleaning workflow can be executed through run_cleaning_pipeline(), which applies the recommended sequence of cleaning operations. Individual functions are also available for users who require a customized workflow; a complete reference to exported functions, arguments, default values, and implementation details is provided in Supplementary Table S1. At the start of each cleaning operation, PhyTaxR ensures that the required cleaning schema is available, allowing atomic use of individual functions without requiring prior execution of the complete pipeline.

The workflow standardizes character encoding and common typographic variants, including quotation marks, dashes, whitespace variants, underscores, and separator characters. It then generates a normalized text representation suitable for subsequent parsing and taxonomic matching in a new column taxon_clean.

#### Extraction of ancillary information

The cleaning pipeline separates the taxonomic core of a record from information that may be biologically informative but can interfere with automated name matching. Extracted information may include taxonomic qualifiers, uncertainty markers, parenthetical annotations, bracketed comments, size ranges, author citations, and comma-separated annotations. These elements are retained in tax_epithet, while uncertain is set when the input indicates an uncertain identification (e.g., cf., aff., or ?).

The workflow also extracts morphological, reproductive, ecological, and descriptive terms, including size classes and descriptors such as centric, pennate, colonial, benthic, or cyst. This approach preserves contextual information from observational records while producing a cleaner taxonomic query string.

#### Taxonomic-core standardization

The remaining taxonomic component is standardized by normalizing infraspecific rank notation, removing residual punctuation where appropriate, and treating short interstitial tokens or abbreviated genus initials. Entries containing alternative or aggregated labels separated by /, &, +, or or can be split into separate records while retaining information on the associated taxon in tax_epithet.

Generic or vernacular group labels are mapped to appropriate higher taxonomic groups where possible. Optional vernacular corrections can also be applied through a persistent user-managed dictionary. Together, these operations transform heterogeneous monitoring labels into taxonomically interpretable inputs for the subsequent automatic-resolution stage.

### Stage 2: Conservative automatic taxonomic resolution

The automatic resolution workflow is executed through run_resolution_pipeline() and is applied to the cleaned taxonomic names stored in taxon_clean. The workflow follows a sequential and conservative strategy: each method is applied only to records unresolved by preceding methods.

`First, search_worms_priority()` performs an exact lookup in WoRMS. For unresolved records, `search_worms_Taxamatch()` performs Taxamatch-based queries to identify close matches, including likely orthographic variants. Remaining unresolved records are evaluated using `search_gbif_strict()`, which accepts GBIF matches only when the API reports an exact match (`matchType = “EXACT“`), a match-confidence score of at least 99 out of 100, and the matched name can be cross-validated against WoRMS. The confidence score is returned by GBIF’s matching algorithm and reflects the strength of the match based on the name and associated taxonomic information. PhyTaxR uses the *≥* 99 threshold as an additional conservative acceptance criterion, alongside exact matching and WoRMS cross-validation. This ordered cascade is designed to maximize the proportion of records resolved automatically while limiting false-positive assignments: progressively less direct matching procedures are applied only after more conservative methods have failed, and GBIF-derived matches are accepted only when they meet strict confidence and cross-validation criteria.

For WoRMS-resolved records, matched_name and matched_aphiaid record the name and AphiaID returned by the matching procedure. `resolve_taxonomic_status()` then determines whether the matched record is currently accepted. When the matched record is a synonym or another non-accepted taxonomic name, the function follows the valid_AphiaID pointer and stores the currently accepted taxon in `accepted_name and accepted_aphiaid`; when the matched record is already accepted, the accepted name is identical to the matched name. For subsequent taxonomy retrieval, `get_taxonomy()` uses accepted_aphiaid when available and otherwise uses matched_aphiaid, thereby retrieving the classification associated with the currently accepted taxonomic concept, including ranks from kingdom to forma.

To improve performance when processing large sets of unresolved taxonomic names, `search_worms_fuzzy_minor()` parallelizes candidate-match evaluation using the future and furrr frameworks. After candidate records have been retrieved from WoRMS, string-similarity calculations and genus-validation procedures are distributed across independent multisession workers through `furrr::future_map2()`. The number of workers is controlled through the ncores argument, which defaults to two workers, and the parallelization plan is reset to sequential execution after processing. External API requests remain rate-limited and are performed separately from the parallel similarity calculations, reducing the likelihood of excessive query rates while preventing temporary service failures from interrupting the entire workflow.

### Stage 3: Semi-automatic fuzzy resolution

Records that remain unresolved after the conservative automatic stage can be processed through the semi-automatic fuzzy-resolution workflow. This stage is intended for ambiguous records, unusual spelling variants, abbreviated names, and labels for which automated acceptance would not provide sufficient confidence. The workflow uses genus and epithet vocabularies constructed from previously resolved records, string-similarity metrics, and fuzzy WoRMS queries to generate candidate matches. Candidate suggestions are presented for user inspection, and the user can accept a proposed match, retain a record for expert review, or flag it for removal. This design ensures that uncertain matches are not silently incorporated into the final dataset. The fuzzy workflow supports checkpointing, allowing a review session to be resumed without repeating prior decisions. User-reviewed decisions can subsequently be encoded in a user-maintained vernacular dictionary, allowing these corrections to be applied consistently in future workflow executions.

### Worked example and reproducibility

A complete worked example is provided in Supplementary Script S1. The script applies the wrapper-based ‘PhyTaxR’ workflow to a small illustrative dataset of raw phytoplankton labels representing common sources of taxonomic ambiguity, including qualifiers, morphological descriptors, size information, generic labels, and potential spelling variants. The dataset is first processed with run_cleaning_pipeline(), optionally followed by vernacular corrections, and then passed to run_resolution_pipeline(). Records that remain unresolved after the automatic stage are processed, where required, through the interactive process_fuzzy_batch() workflow. Supplementary Script S2 applies the same workflow to the same illustrative cases using the individual functions from Stages 1–3 in their execution order. This stepwise presentation makes the transformations applied at each stage explicit and allows to inspect, reproduce, or adapt individual components of the workflow. The behaviour, arguments, and default settings of the exported functions are documented in Supplementary Table S1. To support reproducibility, the supplementary materials also provide the package version, R session information, dates of external database queries, example input data, generated workflow outputs, and the complete sequence of function calls. The outcomes of this worked example are reported in the Results section below.

## Results

Applying the full three-stage workflow (Supplementary Script S1) to 50 raw taxon labels yielded 52 cleaned records after aggregate entries were split into independent rows. Of these, 47 (90.4%) were resolved automatically in Stage 2, and 4 more were resolved through explicit user review in Stage 3, leaving a single record flagged for expert attention. The following subsections describe the outcomes of each stage in turn.

### Stage 1 outcomes: cleaning

The wrapper `run_cleaning_pipeline()` from the first step transformed the 50 raw labels into 52 records: `split_separator_entries()` expanded the two aggregate entries (Chaetoceros decipiens / Skeletonema costatum, C21; Ceratium furca/tripos, C46) into independent records, each retaining a cross-reference to its counterpart in tax_epithet. Across the 52 resulting records, only 34 distinct taxon_clean strings remained, reflecting the deliberate redundancy of the edge-case set, in which several orthographic and annotation variants of the same underlying name (e.g., Chaetoceros decipiens, generated from ten different raw labels including author citations, uncertainty qualifiers, size ranges, and typographical errors) collapse onto a single standardized string (Table 2).

**Table 2.**
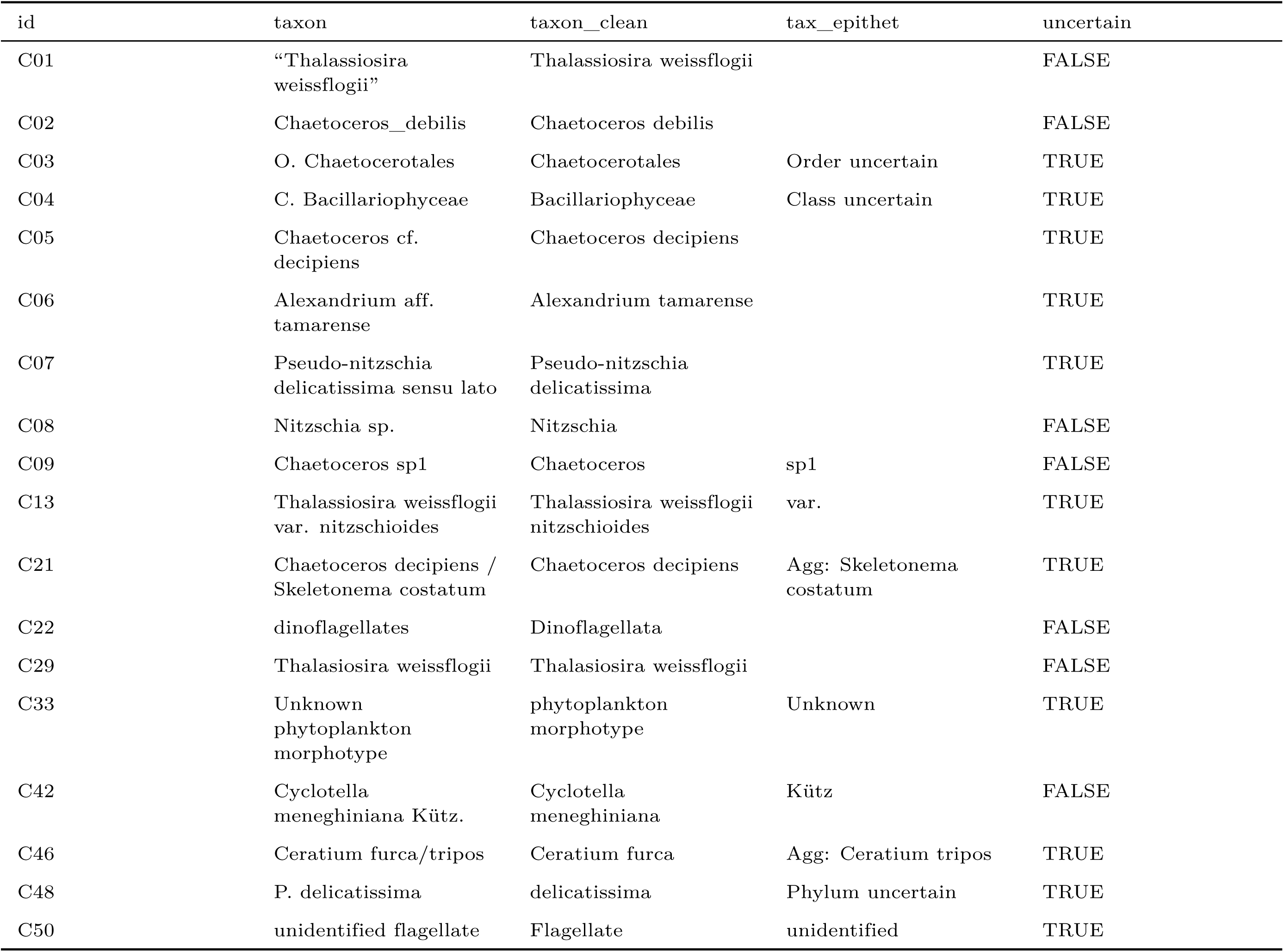
Illustrative outcome of the Stage 1 cleaning pipeline (run_cleaning_pipeline()) for a representative subset of the 52 cleaned records.

The cleaning pipeline consistently separated the taxonomic core from ancillary information regardless of the type of annotation involved: taxonomic-rank prefixes (O. Chaetocerotales, C03), author citations (Cyclotella meneghiniana Kütz., C42), abbreviated genus initials (Chaetoceros ch affinis, C27), and informal infraspecific notation (Thalassiosira weissflogii var. nitzschioides, C13) were all relocated to tax_epithet without discarding the underlying information, while uncertain was set whenever the label carried an explicit qualifier (cf., aff., sensu lato) or an unresolved descriptor (Unknown phytoplankton morphotype, C33). This separation exists because taxonomic backbases such as WoRMS and GBIF match on exact or near-exact scientific names; retaining unprocessed annotations in the query string would systematically depress match rates in Stage 2, while discarding them outright would destroy potentially useful biological context (size class, reproductive stage, or collector confidence) that downstream users may wish to recover.

### Stage 2 outcomes: automatic resolution

`run_resolution_pipeline()` resolved 47 of the 52 cleaned records (90.4%) without requiring user intervention (Table 3): 39 records were matched exactly in WoRMS (worms_exact), 8 were resolved through Taxamatch-based queries (worms_Taxamatch), and none of the 5 candidates passed to search_gbif_strict() met its strict confidence and cross-validation criteria.

**Table 3.**
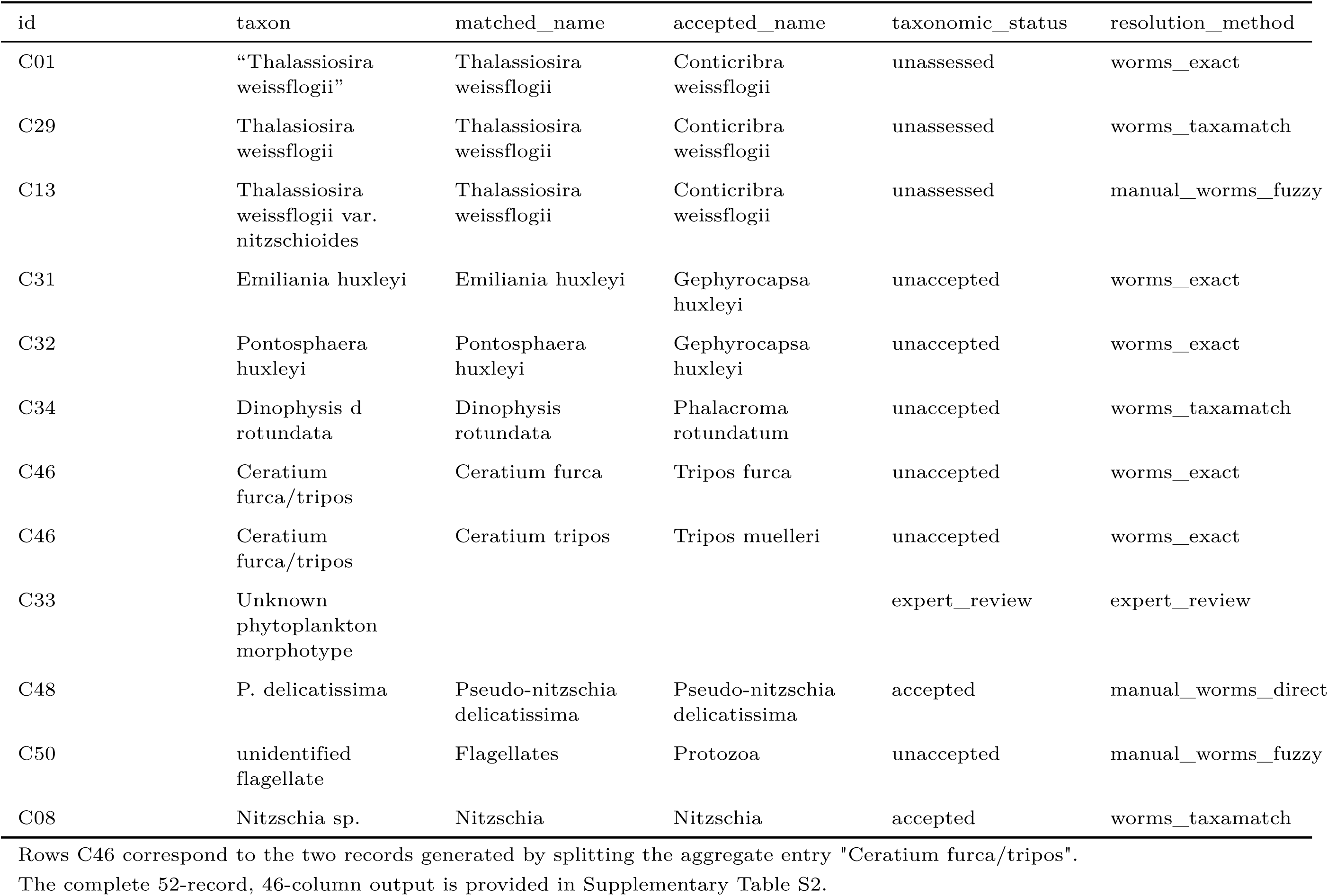
Resolution outcome for a representative subset of the 52 cleaned records, illustrating synonym harmonization, genus reclassification, and the three possible endpoints of the workflow (accepted, unaccepted/synonym, and unresolved/expert review).

The cascade order (exact match, then Taxamatch, then strict GBIF) is conservative by design: each subsequent method is applied only after the preceding one has failed, so that progressively less direct matching procedures are invoked only when necessary, minimizing the opportunity for false-positive assignments. This is illustrated directly by C29 (Thalasiosira weissflogii, a one-letter typographical error), which could not be matched exactly but was correctly resolved via Taxamatch to the same accepted concept as three other noisy variants of the same name (C01, C11, and, after Stage 3, C13) — run_resolution_pipeline() unifies all four into Conticribra weissflogii, the currently accepted replacement for the historical name *Thalassiosira weissflogii*. A second, independent example of this harmonization is provided by C31 (Emiliania huxleyi) and C32 (Pontosphaera huxleyi): two taxonomically unrelated-looking historical names that both resolve, via resolve_taxonomic_status() following the valid_AphiaID pointer, to the single currently accepted taxon *Gephyrocapsa huxleyi*. Without this explicit synonym-resolution step, these records would appear as up to four distinct taxa in a downstream analysis rather than the two biological entities they actually represent — precisely the double-counting risk that motivates the traceability design of PhyTaxR.

For every resolved record, get_taxonomy() retrieves the full WoRMS classification associated with the accepted taxonomic concept (accepted_aphiaid when available, otherwise matched_aphiaid), populating a fixed set of rank-specific columns from kingdom to forma rather than the variable-depth, list-based structure returned natively by wm_classification() in worrms. This design choice serves two purposes consistent with the goals stated in the Introduction: first, retaining minor and intermediate ranks (e.g., Chaetocerotanae, Bacillariophycidae) rather than only the major clades preserves information relevant to fine-grained ecological grouping; second, mapping every record onto the same fixed-width schema, regardless of how many ranks WoRMS assigns to a given taxon, produces a tidy, rectangular output that integrates directly with dplyr/tidyr verbs (filter(), group_by(), pivot_longer()) without requiring users to first flatten a nested taxonomic structure.

### Stage 3 outcomes: semi-automatic fuzzy resolution

The 5 records that remained unresolved after Stage 2 (C13, C30, C33, C48, C50) were passed to process_fuzzy_batch(). Of these, only 3 (C13, C30, C33) were eligible for the automated, non-interactive search_worms_fuzzy_minor() correction attempted earlier within Stage 2, because that guarded step deliberately restricts itself to multi-word candidate strings of at least six characters; single-token queries such as delicatissima (C48) and Flagellate (C50) are excluded by design, since a single truncated or generic token carries too little information for an automatic edit-distance correction to be trusted. None of the 3 eligible candidates met the 0.85 minimum-similarity threshold automatically, so all 5 records were deferred to explicit human review, consistent with the conservative philosophy applied throughout the package.

Reviewer decisions covered the full range of outcomes the semi-automatic stage is designed to support (Table 3). For C30 (Chaetoceros decipens, a one-letter misspelling), the top-ranked, similarity 1.00 candidate (Chaetoceros decipiens) was accepted directly. For C13, the reviewer accepted a “binomial parent” suggestion that resolves the record to the species level with the infraspecific epithet retained as an explanatory note, since no direct record for the variety nitzschioides exists in WoRMS under the accepted genus. For C48 (P. delicatissima), the top-ranked automatic suggestion (Delicatiscala albida, similarity = 0.55) was a taxonomically implausible long shot; rather than accepting it, the reviewer entered the correct scientific name manually, which process_fuzzy_batch() resolved directly to the currently accepted taxon *Pseudo-nitzschia delicatissima* — the same accepted concept independently reached by three other records in the dataset (C07, C25, C37) through ordinary Stage 2 matching, underscoring that manual entry in Stage 3 is a deliberate escape hatch for cases where automatic candidate generation fails, not a lower-confidence alternative to it. For C50 (unidentified flagellate), a “good” candidate (Flagellates, similarity = 0.91) was accepted, but because this candidate is an unaccepted synonym recorded at kingdom rank, the accepted name resolved only to *Protozoa* — illustrating that a high string-similarity score does not guarantee a fine-grained taxonomic resolution when the underlying label is itself generic. Finally, C33 (Unknown phytoplankton morphotype) returned no viable candidates at any stage of the cascading fuzzy search and was explicitly flagged for expert review rather than forced into an arbitrary match; this record therefore remains unresolved (resolution_method = “expert_review”) in the final output, a deliberate design outcome rather than a failure of the workflow.

After Stage 3, 51 of the 52 records (98.1%) in the illustrative dataset were resolved to a currently accepted taxonomic concept, with the single remaining record explicitly and traceably flagged for expert attention rather than silently discarded or force-matched.

## Discussion

PhyTaxR addresses a gap left open by general-purpose taxonomic tools such as taxize and TNRS/RTNRS [10, 11]: rather than assuming a relatively clean scientific name as input, it explicitly handles the noisy, composite structure of phytoplankton monitoring labels, in which a single field may combine a Latin binomial with size ranges, morphological descriptors, uncertainty qualifiers, author citations, and informal group terms. By separating the taxonomic core from ancillary information before external queries are issued (Stage 1), PhyTaxR is designed to increase the proportion of records that can be resolved through direct, conservative matching rather than fuzzy inference, while retaining contextual information that may be useful for subsequent interpretation or review. In the illustrative dataset, 90.4% of cleaned records were resolved automatically through exact or Taxamatch-based WoRMS queries alone, without accepting a GBIF or similarity-based correction. This rate should not be interpreted as a universal expectation, because the example was designed to illustrate common curation challenges rather than to provide a representative benchmark of operational monitoring datasets. Automatic-resolution rates may differ with the composition of the input data, including the prevalence of repeated names, ambiguous labels, and taxa represented in the reference databases.

Beyond the number of records resolved, a central contribution of the package is traceability. Because resolve_taxonomic_status() follows the valid_AphiaID pointer for matched records, PhyTaxR makes explicit when apparently different historical names refer to the same currently accepted taxon. For example, *Emiliania huxleyi* and *Pontosphaera huxleyi* both resolved to *Gephyrocapsa huxleyi*, while the two records derived from the aggregate label *Ceratium furca/tripos* resolved to accepted names in the genus *Tripos*. A workflow that stopped at the first exact match, or discarded synonym information after matching, could reproduce the double-counting and cross-study inconsistency problems that motivate taxonomic harmonisation. In contrast, PhyTaxR retains the matched name, accepted name, identifiers, taxonomic status, resolution method, and explanatory notes together, allowing users to inspect how each decision was reached and to determine how historical identifications should be treated in downstream analyses.

Retrieval of the full WoRMS classification, from kingdom to forma where available, extends this traceability to taxonomic grouping. Rather than returning only a few major clades or a variable-depth nested classification, PhyTaxR stores rank-specific fields in a fixed, tidy schema. This output can be filtered, joined, summarized, or reshaped by any available rank with standard dplyr and tidyr operations, without requiring users to flatten a list-based taxonomy object first. The approach is particularly useful for phytoplankton datasets, where analyses may need to be conducted at several taxonomic resolutions, from broad functional or morphological groups to genera, species, varieties, or formae.

The heterogeneity of real-world taxonomic labels also means that fully unsupervised resolution is neither desirable nor, in practice, achievable. The illustrative dataset included failure modes commonly encountered in monitoring records: typographical errors (Chaetoceros decipens), informal or generic group labels (unidentified flagellate, dinoflagellates), inconsistent infraspecific notation, and outdated genus placements. Five records could not be resolved by the conservative automatic cascade, and Unknown phytoplankton morphotype returned no viable candidate at any stage of fuzzy searching. Rather than forcing an arbitrary assignment, the workflow allows to retain these kind of records and explicitly flag them for expert review. This is a deliberate design choice: an incorrect automatic assignment is generally more damaging to downstream analyses than an unresolved record with documented uncertainty.

Stage 3 complements the conservative cleaning and automatic-resolution stages by providing a structured route for recovering records that would otherwise remain unresolved or be excluded from later analyses. Reviewers may accept well-supported suggestions, enter an authoritative name or AphiaID manually, retain a record for expert review, or flag it for removal. For instance, the low-similarity suggestion generated for P. delicatissima was rejected in favour of a manually entered, taxonomically appropriate name, whereas the one-letter misspelling Chaetoceros decipens was resolved after the reviewer accepted a high-confidence candidate. This distinction is important: fuzzy matching is used to support human judgment rather than replace it. The resulting three-way outcome—resolved automatically, resolved after review, or retained for expert attention—makes residual uncertainty visible and preserves records that may later be resolved using additional ecological, morphological, or regional knowledge.

General-purpose R packages for taxonomic name resolution, including taxize, taxadb, and TNRS/RTNRS, together with direct interfaces to GBIF’s species/match endpoint, provide broad taxonomic and backbone coverage. However, they typically assume that input names are already reasonably well formed and offer limited dedicated support for separating morphological descriptors, size information, uncertainty markers, and informal group labels from the taxonomic core before matching. PhyTaxR instead narrows its scope to marine and estuarine phytoplankton and combines this domain focus with a WoRMS-first, GBIF-secondary cascade, in which GBIF matches are accepted only when they are exact, high-confidence, and independently cross-validated against WoRMS. This design trades taxonomic breadth for precision and domain fit. It is not intended to replace broader taxonomic tools, but to provide a complementary and conservative front end for noisy phytoplankton labels. Its tidy input and output structures, modular functions, and high-level wrappers also allow the package to be incorporated into existing R workflows alongside other tools, without requiring substantial adaptation of data structures or downstream analyses. A direct quantitative comparison of automatic-resolution rates between PhyTaxR and general-purpose tools on the same phytoplankton datasets was outside the scope of this study and remains an important topic for future evaluation.

The package also has limitations. Its automatic-resolution stages depend on the availability, coverage, and ongoing taxonomic maintenance of the WoRMS and GBIF services. Although timeouts, retries, rate limiting, and query-date metadata support robust and reproducible use of these services, changes in external taxonomic backbones or temporary service interruptions remain outside the package’s control. Similarly, records absent from WoRMS, or labels containing insufficient taxonomic information, cannot be resolved automatically with confidence. These limitations reinforce the importance of retaining intermediate outputs, resolution notes, and explicit expert-review flags rather than treating taxonomic resolution as a one-time, irreversible transformation. Furthermore, PhyTaxR does not eliminate the broader fragmentation of software practices in marine science, where researchers often learn tools informally and develop individualized, ad hoc workflows. However, its modular functions, documented high-level workflows, user-managed correction mechanisms, vignette, versioned releases, and reproducible supplementary scripts are intended to reduce barriers to reuse and to make taxonomic-curation decisions more transparent across users and projects [12]. By formalizing a workflow that would otherwise remain embedded in project-specific scripts, the package provides a reusable basis for phytoplankton taxonomic curation while preserving the possibility of expert intervention where automated evidence is insufficient.

## Acknowledgments

We would like to thank Rose Terner and Mohammad Al-Mamun for testing PhyTaxR during its development and for their constructive suggestions to improve its functionality. We also want to thank Mohammad M. Amirian for his advice on the development of the package, and for his helpful comments on the preparation and organization of this manuscript.

## Supporting information

**S1 Script. Wrapper-based phytaxr workflow for the 50-case illustrative dataset.**

**S2 Script. Function-level phytaxr workflow for the 50-case illustrative dataset.**

**S1 Table. Function-level reference for phytaxr version 0.3.2.** Function purposes, inputs, outputs, arguments, default values, and source-code links.

**S2 Table. Record-level cleaning and taxonomic-resolution results for the illustrative dataset.** Curation fields and taxonomic classifications for the 52 records generated from the 50-case input dataset.

**S1 File. Input data, workflow outputs, and query metadata for the illustrative example.** ZIP archive containing the unified input dataset, all files from the data directory, and the complete 52-record, 46-column output in CSV format.

